# Polymorphisms in the vitamin D receptor gene are associated with reduced rate of sputum culture conversion in multidrug-resistant tuberculosis patients in South Africa

**DOI:** 10.1101/132852

**Authors:** Matthew J Magee, Yan V. Sun, James CM Brust, N. Sarita Shah, Yuming Ning, Salim Allana, Angela Campbell, Qin Hui, Koleka Mlisana, Pravi Moodley, Neel R Gandhi

## Abstract

**Background:** Vitamin D modulates the inflammatory and immune response to tuberculosis (TB) and also mediates the induction of the antimicrobial peptide cathelicidin. Deficiency of 25-hydroxyvitamin D and single nucleotide polymorphisms (SNPs) in the vitamin D receptor (VDR) gene may increase the risk of TB disease and decrease culture conversion rates in drug susceptible TB. Whether these VDR SNPs are found in African populations or impact multidrug-resistant (MDR) TB treatment has not been established. We aimed to determine if SNPs in the VDR gene were associated with sputum culture conversion among a cohort of MDR TB patients in South Africa.

**Methods:** We conducted a prospective cohort study of adult MDR TB patients receiving second-line TB treatment in KwaZulu-Natal province. Subjects had monthly sputum cultures performed. In a subset of participants, whole blood samples were obtained for genomic analyses. Genomic DNA was extracted and genotyped with Affymetrix Axiom Pan-African Array. Cox proportional models were used to determine the association between VDR SNPs and rate of culture conversion.

**Results:** Genomic analyses were performed on 91 MDR TB subjects enrolled in the sub-study; 60% were female and median age was 35 years (interquartile range [IQR] 29-42). Smoking was reported by 21% of subjects and most subjects had HIV (80%), were smear negative (57%), and had cavitary disease (55%). Overall, 87 (96%) subjects initially converted cultures to negative, with median time to culture conversion of 57 days (IQR 17-114). Of 121 VDR SNPs examined, 10 were significantly associated (p<0.01) with rate of sputum conversion in multivariable analyses. Each additional risk allele on SNP rs74085240 delayed culture conversion significantly (adjusted hazard ratio 0.30, 95% confidence interval 0.14-0.67).

**Conclusions:** Polymorphisms in the VDR gene were associated with rate of sputum culture conversion in MDR TB patients in this high HIV prevalence setting in South Africa.

**Author contributions:** MJM, YVS, JCMB, SS, and NRG conceived and designed the study and drafted the initial manuscript. MJM, YVS, YN, and QH performed the data analyses. All authors contributed to interpretation of the data, revised the manuscript, and approved the final version.

The findings and conclusions in this article are those of the authors and do not necessarily represent the views of the Centers for Disease Control and Prevention (CDC). The use of trade names and commercial sources is for identification only and does not imply endorsement by the CDC.

## Introduction

In 2015 there were an estimated 480,000 cases of multidrug-resistant (MDR) tuberculosis (TB) worldwide [1]. MDR TB (resistance to at least isoniazid and rifampin) treatment is less effective, more toxic and costly compared to drug susceptible TB [2]. Importantly MDR TB is associated with poor TB treatment outcomes and increased risk of death [3,4], in 2014 there were an estimated 190,000 deaths from MDR TB [5]. Despite global efforts, MDR TB remains difficult to diagnose and treat, and few new therapeutic options are available [6].

Given the paucity of new drugs available for the treatment of MDR TB there has been substantial clinical interest in adjunctive use of 25-hydroxyvitamin D (vitamin D) to improve TB—including MDR TB—treatment outcomes [7]. Vitamin D has anti-inflammatory and anti-bacterial properties that could theoretically improve clinical TB outcomes. The active metabolite of vitamin D, calcitriol, mediates innate immune responses via the induction of the antimicrobial peptide cathelicidin and reactive oxygen intermediates. Vitamin D also promotes macrophage-mediated killing of *Mycobacterium tuberculosis* and modulates both anti-inflammatory and pro-inflammatory T-helper responses to TB [8,9]. Low exposure to solar ultraviolet light, inadequate intake of vitamin D and its precursors, or particular genotypes of the vitamin D receptor (VDR) may lead to vitamin D deficiency which, in turn, could inhibit reduction of bacillary burden, inhibit culture conversion, and impair the effectiveness of TB treatment. Despite the hypothesized plausibility that vitamin D supplementation may improve TB treatment outcomes and the pervasiveness of vitamin D deficiency, clinical trials to date among patients with drug susceptible TB have not demonstrated efficacy in improving rate of sputum culture conversion [10-12]. However, evidence suggests that the effects of vitamin D may vary based on vitamin D receptor (VDR) genotypes, implying that supplementation may only be of clinical benefit in subpopulations with a particular VDR genotype [13].

In certain ethnic populations, single nucleotide polymorphisms (SNPs) in the vitamin D receptor (VDR) gene may increase the risk of TB disease [14]. Three previous studies have reported an association between VDR gene polymorphisms and smear or culture conversion time in patients with pulmonary TB [15-17]. However, the extent to which VDR SNPs are found in African populations or impact MDR TB treatment is limited. South Africa has a high burden of drug-resistant TB, with an estimated 13,000 MDR TB cases in 2014 and 2% MDR TB prevalence rate among all new TB cases [5]. In this context, we aimed to determine if SNPs in the VDR gene were associated with time to sputum culture conversion among a cohort of MDR TB patients in South Africa.

## Methods

### Parent Study

The SHOUT study was a prospective observational cohort study among patients receiving second-line TB treatment for MDR TB from three sites in KwaZulu-Natal province, South Africa between 2011 and 2015. KwaZulu-Natal is the South African province that has been most severely affected by TB (incidence: 1076 cases per 100,000 population) and HIV (prevalence: 17%) [18,19]. Patients 18 years or older were eligible to participate if they had a sputum culture positive for *M. tuberculosis* with phenotypic resistance to both isoniazid and rifampin. Patients with unknown HIV status at the time of enrollment were offered an HIV test. Patients were excluded if they had previous MDR TB treatment, resistance to either fluoroquinolones or injectable TB medications, renal or hepatic dysfunction, or were pregnant. Subjects were treated with the standardized South African MDR TB regimen of kanamycin, moxifloxacin, ethionamide, terizidone, ethambutol, and pyrazinamide. Kanamycin was typically given for a minimum of 6 months, or 4 months after culture-conversion, and oral medications were continued without kanamycin for an additional 12-18 months after culture-conversion. All HIV co-infected participants were initiated on ART within 2 months of MDR TB treatment initiation, regardless of CD4 count, if they were not already receiving ART. Standard ART regimens consisted of efavirenz, and stavudine and lamivudine prior to 2013, or tenofovir and emtricitabine afterwards. Study participants were seen monthly for follow-up for the duration of MDR TB treatment, which is typically 21-24 months.

### Study population, Measures and Definitions

The current study was conducted among a subset of patients from the parent SHOUT MDR TB study who consented to have whole blood samples collected for genomic analyses. Sub-study patients had the same inclusion/exclusion criteria as the parent study and were enrolled beginning in April 2013 until the target (n=100) was reached.

### Measures and Definitions

The primary outcome of interest for the study was time to initial sputum culture conversion. Sputum cultures were performed monthly during MDR TB treatment. Mycobacterial cultures and drug-susceptibility testing (DST) were performed as previously described [4]. Time to sputum culture conversion was defined by the number of days between MDR TB treatment initiation and the first of two negative sputum cultures [20]. Patients who converted sputum cultures to negative before the start of MDR TB treatment were defined as having a 1 day conversion time. A secondary outcome of interest was poor tuberculosis treatment outcome and was defined as a patient who died, interrupted treatment, or had treatment failure.

The primary exposures of interest were VDR gene polymorphisms. Participant samples were genotyped using the Affymetrix® Axiom Pan-African Array according to the manufacturer’s instructions [21]. Quality checks were performed to ensure the overall SNP call rate was >95% and that there was no sex mismatch between genotypic and phenotypic measurement. SNPs were excluded if they had an unknown chromosomal location, a call rate less than 95%, a Hardy-Weinberg Equilibrium (HWE) *p*-value less than 0.0001 or a minor allele frequency (MAF) less than 0.05. After quality control filters, 1,494,763 SNPs were available for genetic analysis. South African samples were pooled with HapMap EUR, YRI and ASW populations to identify population structure relative to European and West African ancestry [22]. Top principal components (PCs) were calculated using independent SNPs after pruning by pair-wise linkage disequilibrium R^2^ larger than 0.1 within windows of 50 SNPs. Using base pair location of human genome build 37, there were 121 SNPs annotated to the VDR gene and passed quality control filters.

Smoking status and alcohol use were self-reported by study patients. Both smoking status and alcohol use were categorized dichotomously as current use (yes/no).

### Statistical Analyses

We examined 121 VDR SNPs to assess their association with rate of sputum culture conversion among patients with MDR TB. Each SNP was coded using the additive genetic effect. Cox proportional models were used to determine the association between VDR SNPs (additive effect) and the hazard rate of initial sputum culture conversion. Patients who had a positive culture at time of MDR TB diagnosis but converted sputum cultures to negative before the start of MDR TB treatment were censored at day 1. Patients who died or failed treatment before an initial sputum culture conversion were censored. All models were adjusted for age, sex, smoking status, alcohol, AFB smear status, HIV status, and cavitary disease. For the secondary outcome of TB treatment result, we used logistic regression to estimate the odds of poor TB treatment outcome among a subset of SNPs associated with reduced rate of culture conversion. Patients who withdrew from the study before treatment completion were excluded from the secondary outcome analysis.

### Ethics

The study protocol was approved by the institutional review boards at the University of KwaZulu-Natal, Albert Einstein College of Medicine, and Emory University, and by the KwaZulu-Natal Department of Health and CDC’s National Center for HIV, Hepatitis, STDs and Tuberculosis. All participants signed written informed consent.

## Results

From 2011-2013, the parent cohort study screened 365 patients and enrolled 206. Among patients enrolled in the parent study, 103 provided samples for the present sub-study, 7 were not processed due to DNA extraction errors, and 5 patients were late excluded from the parent study due to second line drug resistance. The 91 remaining participants were included in the current sub-study for genomic analyses and of these, 55 (60%) were female and the median age was 35 years (interquartile range [IQR] 29-(Table 1). Most patients were HIV co-infected (n=73, 80%), had undetectable viral load (n=23/42, 55%) and the median baseline CD4 count of 199 cells/mm^3^ (IQR 143-289). Thirty-nine (43%) patients were AFB smear positive, 50 (55%) had cavitary disease, and 74 (81%) had previously been treated for TB. Smoking was reported by 21% of patients.

**Table 1.** Baseline participant characteristics and 2-month sputum culture status.

abbreviations: IQR-interquartile range; AFB-acid fast bacilli; ARV-antiretroviral
| Characteristic | Total<br>N=91<br>N (%) | Sputum culture<br>negative at 2-months<br>N=50 (55.0)<br>N (%) | Sputum culture<br>positive at 2-months<br>N=41 (45.1)<br>N (%) | P<br>value<sup>A</sup> |
| --- | --- | --- | --- | --- |
| Male | 36 (39.6) | 15 (30.0) | 21 (51.2) | 0.04 |
| Median age, years (IQR) | 35 (29-42) | 35 (27-41) | 38 (32-42) | 0.12 |
| Current smoker<sup>B</sup> | 19 (20.9) | 7 (14.0) | 12 (29.3) | 0.07 |
| Alcohol<sup>B</sup> | 28 (30.7) | 14 (28.0) | 14 (34.2) | 0.53 |
| Baseline AFB positive | 39 (42.9) | 18 (36.0) | 21 (51.2) | 0.14 |
| Baseline cavity | 50 (55.0) | 26 (52.0) | 24 (58.5) | 0.53 |
| Median baseline BMI<br>(IQR) N=82 | 21.6 (18.5-24.6) | 21.8 (19.4-24.4) | 21.3 (16.8-24.9) | 0.42 |
| Previous TB treatment | 74 (81.3) | 40 (80.0) | 34 (82.9) | 0.72 |
| HIV seropositive | 73 (80.2) | 40 (80.0) | 33 (80.5) | 0.95 |
| Median baseline CD4<br>(IQR) N=46<sup>C</sup> | 199 (143-289) | 185 (125-266) | 221 (143-309) | 0.61 |
| On ARV at baseline<sup>C</sup> N=57 | 44 (75.7) | 24 (75.0) | 20 (76.9) | 0.66 |
| Median baseline viral load<br>(IQR) N=42<sup>C</sup> | 83 (39-13000) | 65 (39-75278) | 100 (39-3200) | 0.94 |
| Undetectable viral load<br>N=42<sup>C</sup> | 23 (54.8) | 13 (56.5) | 10 (52.6) | 0.80 |

A. 2-side chi-square p-value, except for age (2-sided Wilcoxon rank sum)
B. Self-reported
C. Among HIV positive only

All samples passed standard genome-wide association study (GWAS) quality control filters with the lowest individual level SNP call rate of 98.2%. Using top two PCs from GWAS data (Figure 1), we observed the separation of South Africans from populations with known African ancestry (YRI and ASW), and European ancestry (EUR). PC1 clustered European versus African ancestry, while PC2 further distinguished West Africans and South Africans. Within South African samples in this study, no outlier (3SD) was observed using top 10 PCs, indicating a genetically homogeneous population.

**Figure 1.**
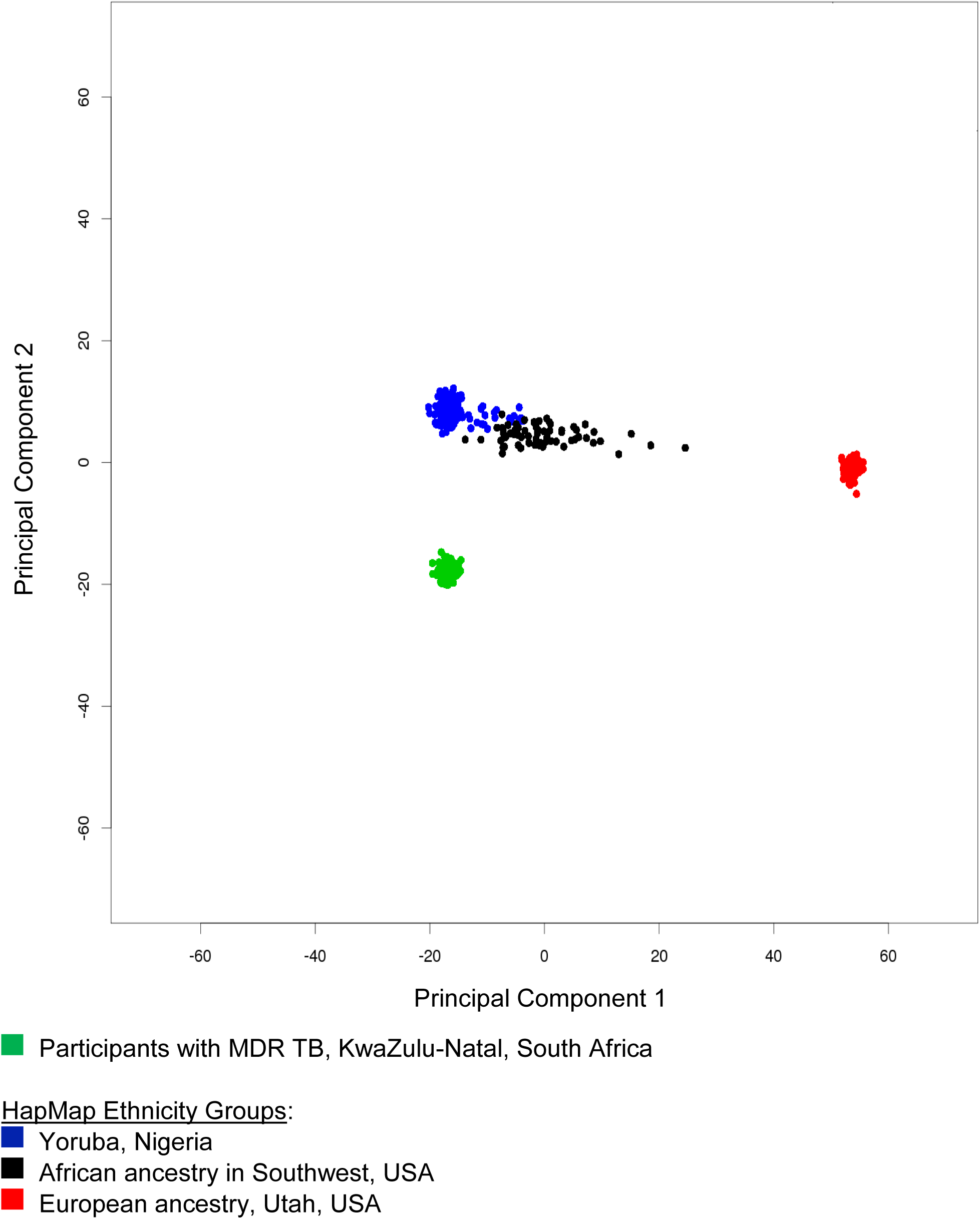
Principal component analysis of study participants compared to HapMap ethnic groups.

Overall, 87 (96%) subjects converted sputum cultures to negative. Of the 87 who converted sputum cultures to negative, 24% (21/87) were positive at the time of MDR TB diagnosis but converted to negative before the time of MDR TB treatment start. The median time to culture conversion was 57 days (IQR 17-114); among patients who were not culture negative at time of treatment initiation the median time to conversion was 82 days (IQR 53-143). Among patients who converted, 50 (55.0%) were culture negative by two months of second-line treatment (Table 1). Compared to females, males were significantly more likely to be sputum culture positive at two months (36.4% [20/55] vs. 58.3% [21/36], *p*=0.04).

Of 121 VDR SNPs examined, 10 were significantly associated (p<0.05) with hazard rate of sputum culture conversion in multivariable analyses (Table 2). The estimated slower conversion rate (adjusted hazard ratio of sputum culture conversion <1.0) ranged from 0.30 (95% CI 0.14-0.67) for rs74085240 to 0.64 (95%CI 0.42-0.98) for rs11168287. For example, each additional risk allele on SNP rs74085240 delayed the rate of culture conversion by 70% (aHR 0.55, 95% CI 0.36-0.85). Two VDR SNPs (rs11168327 and rs11574143) were associated with significantly improved rate (adjusted hazard ratio >1.0) of culture conversion (Table 2).

**Table 2.**
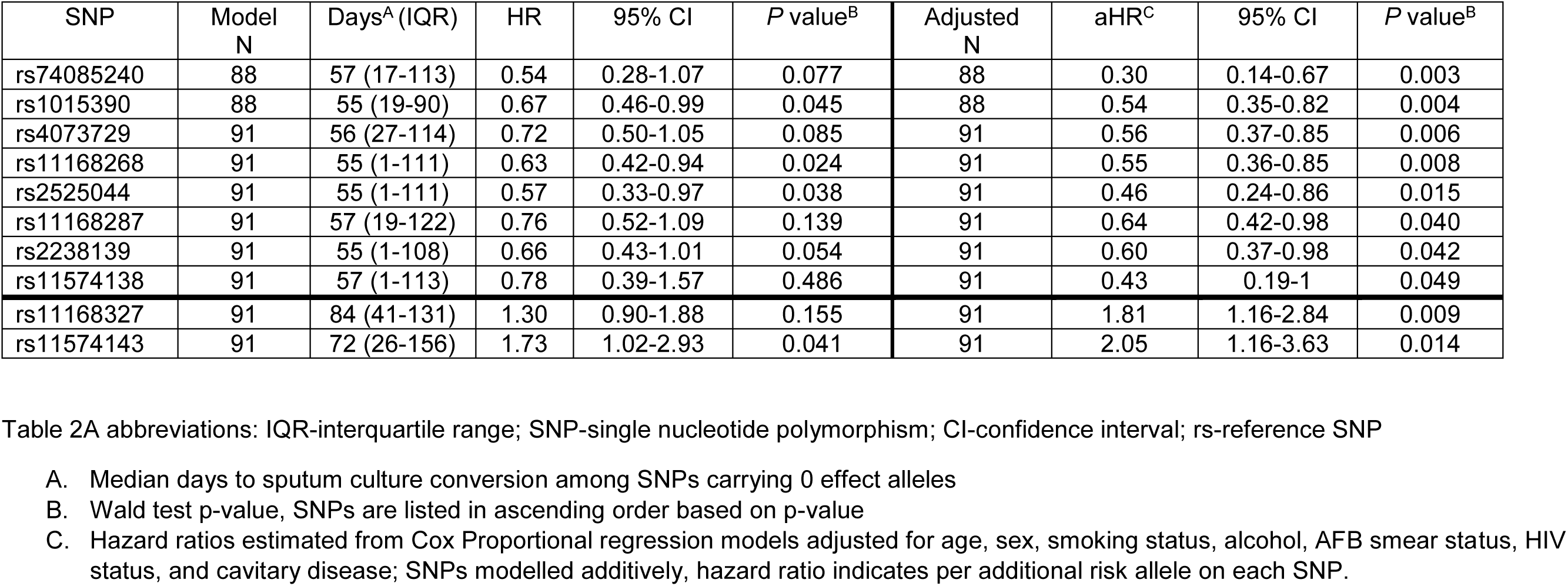
Hazard of sputum culture conversion by vitamin D receptor gene single nucleotide polymorphism.

Overall, 19% (17/88) of patients had a poor TB treatment outcome and 3 additional patients withdrew treatment. We did not detect any genotypes associated with a significant increased odds of poor TB treatment outcome (Table 3).

**Table 3.** Poor tuberculosis treatment outcome by vitamin D receptor gene single nucleotide polymorphism, N=88.

abbreviations: SNP-single nucleotide polymorphism; rs-reference SNP; CI-confidence interval; NA-snp information not available
| SNP | Total<br>N=88 | Poor Outcome<sup>A</sup><br>N=17 (19.3)<br>N (%) | Cured/Completed<br>N=71 (80.7)<br>N (%) | Odds ratio<br>(95% CI) |
| --- | --- | --- | --- | --- |
| rs74085240* |  |  |  |  |
| CC | 2 | 1 (50.0) | 1 (50.0) | 7.50 (0.24-68.83) |
| CA | 8 | 1 (12.5) | 7 (87.5) | 0.58 (0.07-50.09) |
| AA | 76 | 15 (19.7) | 61 (80.3) | 1.00 |
| NA | 2 | 0 | 2 (100) |  |
| rs11574138* |  |  |  |  |
| CC | 1 | 0 | 1 (100) | NA |
| CT | 8 | 3 (37.5) | 5 (62.5) | 2.79 (0.60-13.04) |
| TT | 79 | 14 (17.7) | 65 (82.3) | 1.00 |
| rs11168268* |  |  |  |  |
| GG | 9 | 3 (33.3) | 6 (66.7) | 2.50 (0.50-12.46) |
| GA | 37 | 7 (18.9) | 30 (81.2) | 1.17 (0.37-3.71) |
| AA | 42 | 7 (16.7) | 35 (83.3) | 1.00 |
| rs2525044* |  |  |  |  |
| AA | 2 | 1 (50.0) | 1 (50.0) | 5.09 (0.30-87.68) |
| AG | 19 | 5 (26.3) | 14 (73.7) | 1.82 (0.54-6.09) |
| GG | 67 | 11 (16.4) | 56 (83.6) | 1.00 |
| rs1015390* |  |  |  |  |
| TT | 9 | 2 (22.2) | 7 (77.8) | 0.86 (0.15-1.53) |
| TC | 44 | 6 (13.6) | 38 (86.4) | 0.47 (0.15-5.00) |
| CC | 32 | 8 (25.0) | 245 (75.0) | 1.00 |
| NA | 3 | 1 (33.3) | 2 (66.7) |  |
| rs2238139 |  |  |  |  |
| GG | 5 | 2 (40.0) | 3 (60.0) | 3.33 (0.49-22.60) |
| GA | 23 | 5 (21.7) | 18 (78.3) | 1.39 (0.42-4.62) |
| AA | 60 | 10 (16.7) | 50 (83.3) | 1.00 |

**A.** Poor outcome defined as death or failure.

## Discussion

We examined 121 SNPs in the VDR gene region and found a subset to be associated with rate of sputum culture conversion among patients with MDR TB in South Africa. Specifically, we identified 9 VDR SNPs that were associated with an estimated 50% to 25% reduced (delayed conversion) rate of sputum culture conversion among patients receiving second-line TB therapy. Our findings provide new data about the relationship between VDR polymorphisms and sputum culture conversion among patients with MDR TB in South Africa.

Our findings in patients with MDR TB are consistent with three previous studies that reported significantly lower rates of sputum culture conversion among patients with drug susceptible pulmonary TB who had specific VDR polymorphisms [10,15,17]. First, an observational longitudinal study among 78 Peruvian patients with confirmed pulmonary TB reported that patients with the *TT TaqI* genotype (previous nomenclature now typically replaced by endonuclease digestion pattern) had significantly longer time to sputum culture conversion (median 46 days for *TT* genotype vs. 16 days for *Tt* genotype) [15]. Second, in 2011 Martineau et al conducted a randomized control trial in London, testing the efficacy of 2.5mg vitamin D_3_ supplementation to reduce culture conversion time in smear positive TB patients [10]. The trial reported no overall effect of supplementation on culture conversion rates, but the authors did report an interaction between vitamin D_3_ supplementation and culture conversion with *TaqI* genotype. Specifically, supplementation was beneficial among patients with *tt* genotype (HR 8.09, 95%CI 1.36-48.01). Unlike the studies from Peru and London, we did not observe an association between rs731236 (*TaqI* genotype defined by endonuclease digestion pattern) and rate of sputum culture conversion. Third, an observational longitudinal study of HIV-negative patients with pulmonary TB from the Western Cape of South Africa reported that a significantly lower proportion of patients with *ApaI aa* genotype had converted cultures by month 2 of TB treatment compared to patients with *ApaI Aa* genotype (26% vs. 51%, p=0.03) [17]. All three studies are consistent with our findings that VDR polymorphisms are associated with rate of culture conversion. Unlike the previous studies, our study identified SNPs significantly associated with rate of culture conversion in patients with MDR TB.

To our knowledge, only one previous study examined the association between VDR genotypes and time to culture conversion in patients with MDR TB. In 2012 Rathored et al. followed 236 HIV-negative patients with MDR TB during DOTS-Plus treatment in India and reported no significant differences in bivariate analyses between the three VDR genotypes examined (*BsmI, TaqI, FokI*) and time to culture conversion [16]. However, in our study we examined 121 specific SNPs on the VDR gene region and adjusted for important confounding factors (i.e., smoking status); Rathored et al. did not adjust for confounding which may partially explain why we reported significant differences in rates of culture conversion and the previous study did not. Similar to the study in India, we did not observe a significant association between *BsmI* (rs1544410) or *TaqI* (rs731236) genotypes and time to culture conversion.

Several hypothesized biologic mechanisms may explain why polymorphisms in the VDR gene are associated with rate of sputum culture conversion in patients with MDR TB. Although vitamin D supplementation has not been demonstrated to be efficacious in improving culture conversion time in drug-susceptible TB treatment [10-12], vitamin D likely has a role in TB treatment-induced modulation of circulating immunologic signals. Circulating immune signals are affected by TB treatment alone, for example interleukin (IL)-10, cathelicidin LL-37, and neutrophil gelatinase-associated lipocalin (NGAL) are suppressed by medications administered during the intensive phase TB treatment. Moreover, a randomized trial from London demonstrated that vitamin D supplementation enhanced the immune effects of TB treatment. In the trial of 126 smear-positive pulmonary TB patients, Coussens et al. reported that patients receiving vitamin D supplementation during first-line treatment had TB treatment-induced increases in lymphocytes and reduced concentrations of inflammatory markers [23]. Therefore, it is plausible that effects of second-line TB treatment on the immune responses may be modified differently by vitamin D compared to first-line TB treatment immune modulation by vitamin D.

Our study was subject to limitations. First, we did not measure plasma levels of vitamin D or calcitriol. Therefore, we were unable to verify if vitamin D levels were affected by polymorphisms in the VDR SNPs or if the polymorphisms affected culture conversion directly. Second, we did not measure any immune modulating signals. Vitamin D is hypothesized to affect sputum culture conversion through immune modulation of cytokines (interferon-gamma, IL-2, IL-12) chemokines (chemokine ligand (CXCL)-9, CXCL-10, matrix metallopeptidase-9) and antigen stimulated responses (Th1) [23]. Consequently, we were unable to determine if VDR SNPs influenced the expression of immune modulating signals that may affect rate of culture conversion. Third, our sample had relatively low power and we were therefore unable to adjust statistical tests for multiple comparisons. We did not have power to adjust for covariates in the logistic regression models that were used to estimate the odds of poor MDR TB treatment outcome by VDR SNPs. Fourth, we did not examine sputum culture reversions to positive. The analysis only analyzed the association between VDR SNPs and the patients’ first sputum culture conversion and therefore does not assess the association with sustained conversion.

Despite limitations, our study had several strengths. Foremost, we examined 121 specific SNPs in the VDR gene region among patients with MDR TB from a genetically distinct population. Previous similar studies have focused on VDR polymorphisms at the level of *FokI*, *ApaI*, and *TaqI* genotypes but did not measure specific SNPs and only one previous study enrolled patients with TB from South Africa (which included only drug-susceptible patients). Second, our study only included patients with culture- and DST-confirmed MDR TB and followed patients monthly to obtain cultures during second-line TB treatment. Previous studies examining the association between VDR polymorphisms and response to TB treatment were primarily among drug susceptible patients, included patients without culture confirmed TB, largely examined sputum smear conversion at one time point, and few adjusted for key confounders [24,25].

## Acknowledgements

We are grateful to the study team at the University of KwaZulu-Natal for their tireless efforts in data collection, record abstraction, participant recruitment and interviews. Finally, we thank the participants and their families who consented to participate in this study.

---

**Supplemental Table 1.** Vitamin D receptor gene single nucleotide polymorphisms not significantly associated with initial time to sputum culture conversion

| SNP | P-value<sup>A</sup> | SNP | P-value<sup>A</sup> | SNP | P-value<sup>A</sup> |
| --- | --- | --- | --- | --- | --- |
| rs10875705 | 0.052 | rs7963776 | 0.337 | rs2525049 | 0.634 |
| rs987849 | 0.062 | rs11568820 | 0.341 | rs11574110 | 0.668 |
| rs2239185 | 0.063 | rs2239179 | 0.354 | rs7302235 | 0.671 |
| rs35609792 | 0.070 | rs11168309 | 0.359 | rs60556433 | 0.676 |
| rs10875700 | 0.079 | rs11168328 | 0.366 | rs58187695 | 0.683 |
| rs74088704 | 0.087 | rs11574070 | 0.387 | rs58426141 | 0.718 |
| rs7309452 | 0.092 | rs61919101 | 0.399 | rs4237855 | 0.725 |
| rs11168319 | 0.101 | rs6580642 | 0.401 | rs10467099 | 0.732 |
| rs74086592 | 0.104 | rs7976091 | 0.416 | rs58436504 | 0.735 |
| rs7974905 | 0.110 | rs739837 | 0.419 | rs3819545 | 0.737 |
| rs2246001 | 0.118 | rs12313208 | 0.421 | rs12308082 | 0.740 |
| rs1544410 | 0.120 | rs12717991 | 0.425 | rs10747526 | 0.772 |
| rs4334089 | 0.143 | rs12721416 | 0.428 | rs11168307 | 0.773 |
| rs58379944 | 0.144 | rs2228572 | 0.431 | rs2107301 | 0.778 |
| rs7965281 | 0.158 | rs58789572 | 0.438 | rs11168263 | 0.780 |
| rs11168280 | 0.177 | rs7965943 | 0.444 | rs11574100 | 0.787 |
| rs4442605 | 0.180 | rs11168306 | 0.449 | rs7974708 | 0.796 |
| rs2525043 | 0.183 | rs11574050 | 0.455 | rs10747524 | 0.798 |
| rs757555 | 0.183 | rs7305032 | 0.465 | rs7311030 | 0.799 |
| rs1859281 | 0.186 | rs12314197 | 0.471 | rs2544037 | 0.802 |
| rs4237856 | 0.216 | rs7965274 | 0.475 | rs2239182 | 0.819 |
| rs4393380 | 0.221 | rs11574053 | 0.483 | rs2544039 | 0.834 |
| rs11574044 | 0.222 | rs73109883 | 0.506 | rs10783221 | 0.839 |
| rs2525045 | 0.225 | rs10459227 | 0.515 | rs12721397 | 0.847 |
| rs886441 | 0.241 | rs7970376 | 0.517 | rs12303561 | 0.848 |
| rs2238136 | 0.243 | rs11168325 | 0.538 | rs10459217 | 0.850 |
| rs4254129 | 0.246 | rs7967673 | 0.542 | rs4760648 | 0.856 |
| rs11574041 | 0.252 | rs11574081 | 0.548 | rs12299534 | 0.863 |
| rs7975128 | 0.255 | rs731236 | 0.549 | rs11168261 | 0.877 |
| rs12721370 | 0.270 | rs61553170 | 0.555 | rs61558228 | 0.882 |
| rs11574005 | 0.273 | rs12321826 | 0.558 | rs74085273 | 0.898 |
| rs2238140 | 0.278 | rs11168314 | 0.563 | rs11168264 | 0.910 |
| rs2853560 | 0.291 | rs4341603 | 0.565 | rs2239186 | 0.925 |
| rs4307774 | 0.292 | rs2238138 | 0.577 | rs2189480 | 0.934 |
| rs11168311 | 0.308 | rs2408876 | 0.580 | rs11168277 | 0.953 |
| rs4760674 | 0.331 | rs2544038 | 0.580 | rs3890734 | 0.954 |
| rs4328263 | 0.334 | rs2853561 | 0.619 | rs4760658 | 0.954 |

A. Wald test p-value from Cox Proportional regression models adjusted for age, sex, smoking status, alcohol, AFB smear status, HIV status, and cavitory disease; SNPs modelled additively.

## Author contributions

MJM, YVS, JCMB, SS, and NRG conceived and designed the study and drafted the initial manuscript. MJM, YVS, YN, and QH performed the data analyses. All authors contributed to interpretation of the data, revised the manuscript, and approved the final version.

